# Matrix stiffness induces heritable changes in chromosome numbers, consistent with solid tumor heterogeneity

**DOI:** 10.1101/2025.01.28.635370

**Authors:** Alişya A. Anlaş, Markus T. Sprenger, Mai Wang, Nick Ontko, Steven Phan, Dennis E. Discher

## Abstract

Solid tumors often have an abundance of collagen-I that stiffens the tissue, and they are invariably driven by mutations that include chromosome losses and gains. These observations are linked here by showing that 3D matrix stiffness induces heritable changes to a cell**’**s DNA. We use live-cell chromosome reporters (ChReporters) and hydrogels of tunable stiffness to show mitotic compression, micronuclei counts, ChReporter losses and heterogeneity all increase as functions of stiffness. Increased mistakes occur despite suppressed cell division in stiff matrix and minimal size variation between spheroids. Colonies of ChReporter-negative cells within cancer spheroids align with Luria-Delbruck**’**s seminal theory for heritable mutations, which predicts inter-spheroid variances that exceed Poisson statistics. Suppression of the contractility motor Myosin-II also increases chromosome loss in 3D but not 2D and does not affect spheroid growth – thus clarifying Myosin-II**’**s putative role as a tumor suppressor. Consistent with experiments, pan-cancer analyses of clinical data associates chromosome losses and gains with collagen-I levels and genetic variation. Stiff extracellular matrix thus drives mechano-evolution of solid tumors as a Darwin-Lamarck process with heterogeneity that complicates therapy.

## Introduction

Solid tissues and tumors have an abundance of collagen-I that stiffens extracellular matrix (ECM) and impacts proliferation as well as phenotypic differentiation ^1–6^. A seemingly distinct, additional observation for cancer cells is that errors in division induce losses and gains of multiple chromosomes, which fuels selection for tumor growth ^7–10^. Prevalent in solid tumors moreso than liquid cancers, such genetic heterogeneity complicates many therapies and motivates an understanding of causal factors. Intriguingly, the fraction of the genome with chromosomal losses and gains in well-established human tumors correlates with collagen-I levels for a given cancer type in The Cancer Genome Atlas (TCGA), (**Table 1**, *p* < 0.0001). However, any causal role for ECM or its stiffness in promoting such mutational events remains unclear.

**Table 1.**
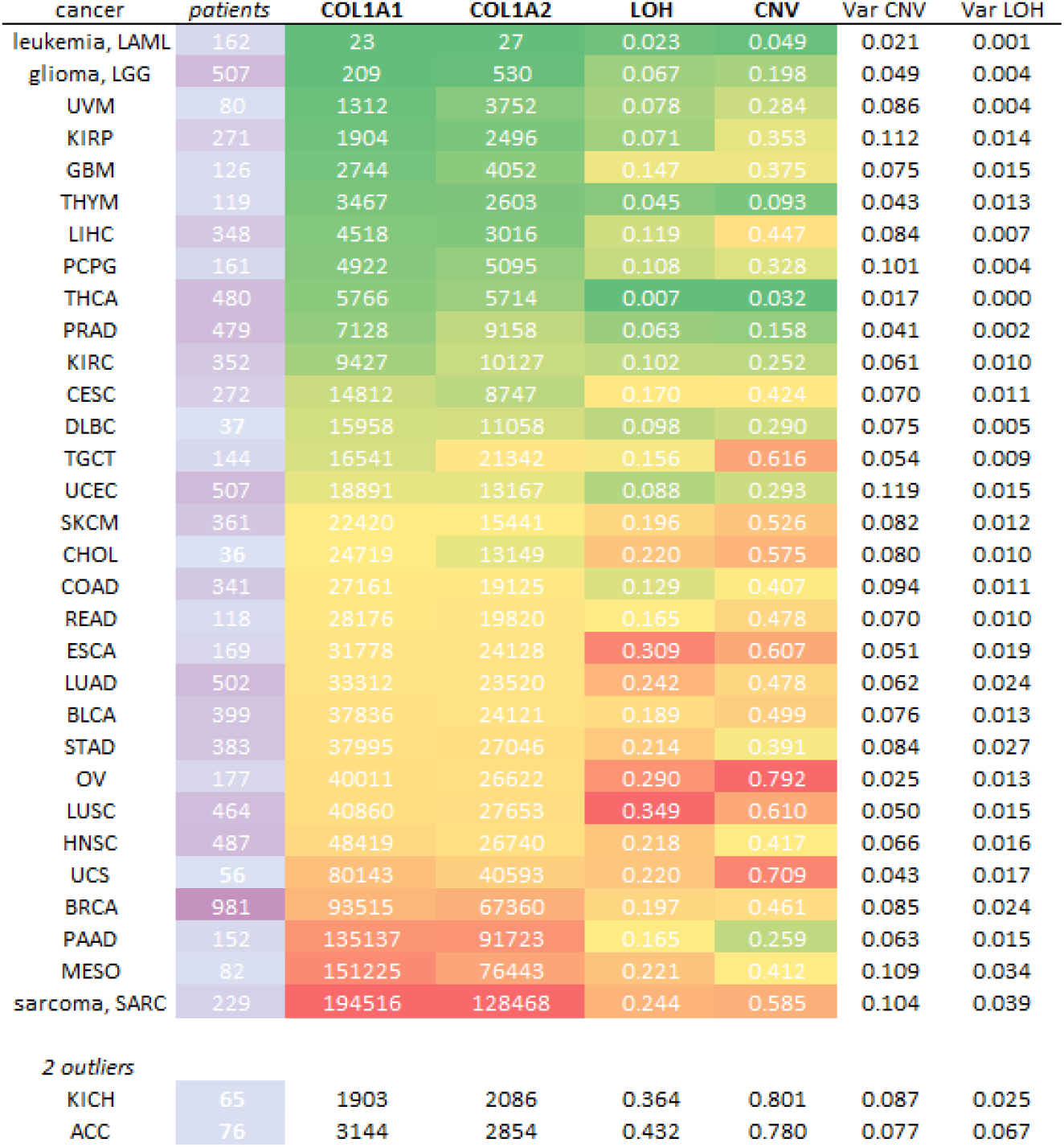
Collagen-1 transcript levels correlate with average chromosome losses or gains across tumor types in The Cancer Genome Atlas (TCGA) (*p* < 0.0001). Standard acronyms for each tumor type are followed by patient numbers, median levels of *COL1* transcripts across patients, and average fractions of the genome showing Loss of Heterozygosity (LOH) or Copy Number Variation (CNV) based on Wang et al. (2023). Interpatient Variance is also listed. All cancer types are included in the Pearson correlation, but two outlier cancer types (KICH, ACC) with low patient numbers are not included in the heatmap scale.

Substrate stiffness in 2D increases cytokinesis failure and can lead to whole genome doubling ^11^ or chromatin bridge fragmentation ^12^. However, ECM stiffness in 3D spheroids and tissue models impedes cell cycle ^4,13,14^ – with cells appearing more confined and compressed ^13,15–19^. Given such mechanical stresses and the noted trends in chromosomal losses and gains (**Table 1**), we hypothesized that rare cell divisions in 3D stiff ECM significantly increase the induction of heritable chromosome losses. We further hypothesized a large variation caused by rare random mutational events *followed by* proliferation in accord with Luria & Delbruck**’**s theory for bacteria^20^ and crowded microbes ^21^. Tests on tumor mutations appear lacking.

Cell stresses can of course cause cell cycle arrest or death, and so induction of rare (<1%) but heritable mutations driven by 3D ECM stiffness is not obvious nor clear. Cell cycle arrest in 2D that results from drug-induced abnormal divisions and aneuploidy is also somehow overcome in 3D spheroids ^22^. Spheroids in viscous methylcellulose media nonetheless grow faster upon knockdown of some known tumor suppressor genes compared to 2D culture, and knockdown of non-muscle myosin-IIA had the same effect ^23^. Myosin-II**’**s roles in multiple mechanosensitive pathways are clear ^1–3,24,25^, and myosin-IIA**’**s knockdown has also been described as causing tumors in mouse epithelia ^26,27^. Mechanisms differ between studies, but any *direct* impact of suppressing myosin-IIA on genetic aberrations in tumor progression remains unclear.

Coupling of 3D tumor models to appropriate visualization methods remains broadly underutilized ^28^– especially for tracking induction of genetic changes. We visualize chromosome losses within live cells that initiate colony formation by gene-editing GFP into a constitutively expressed gene on one chromosome (e.g. GFP-LaminB1 on chromosome 5) (**Fig.1A**) ^16^. We focus on chromosome losses because losses precede gains in some solid tumors ^28^, and our recent studies with **‘**ChReporters**’** in 2D culture models showed easy-to-see chromosome losses as a result of physical or chemical perturbations (i.e. inhibiting the spindle assembly checkpoint with MPS1i). These associated with (i) abnormal mitosis including chromosome mis-segregation in multipolar mitoses and micronucleus formation ^16,29^ linked to complex genome rearrangements ^30,31^. They also associated with (ii) cell cycle delays and cell death. Here, cancer spheroids are cultured within 3D hydrogels that have a stiffness which ranges from that of normal tissue to that of rigid fibrotic or tumor tissues (**Fig.1B**,**C**) ^14,33^. We use agarose because, unlike collagen-I or other native ECM, agarose lacks adhesive ligand; hence, stiffness is the main parameter being tuned by agarose density. As hypothesized, we find stiff 3D microenvironments suppress cell division but do not eliminate it, and heritable chromosome losses as well as mitotic aberrations increase strongly with ECM stiffness. Chromosome loss rates in cancer spheroids not only exceed those in standard 2D cultures but also increase with myosin-IIA disruption. Tumor lines studied are compromised (human lung H23) or normal (mouse melanoma B16) with respect to p53 – as another key **‘**protector**’** of genome integrity.

Importantly, the variance of mutated colonies within and between tumor spheroids extends the threory of Luria and Delbruck ^20,21^ to mechano-evolution as a combined Darwin-Lamarck process where mutations are *stress-enhanced* and *ongoing* per Lamarck. Variations are neither random Gaussian nor Poisson and are not chromosome specific per the determinism of Lamarck. Surprisingly, pan-cancer analyses also show the genetic variance between patients track with mean chromosome changes as do collagen levels. Our results thus offer some explanation as to how solid tumors become genetically heterogeneous.

## Results

### Matrix stiffness in 3D increases division errors

Spheroids of H23 cells in methylcellulose double more slowly than in 2D (∼60 hr vs ∼40 hr: **Figure 1B**), which aligns with more contacts and inhibition in 3D. The spheroid core indeed showed fewer divisions than the surface based on staining for microtubules and DNA (**Figure 1D**). Dividing cells in the core also show ∼20% smaller area compared to those on the surface, which round up into the surrounding fluid (**Figure 1D**). Apoptosis was not prominent in spheroids despite the suggested crowding by neighboring cells (not shown). To measure this cell-confining stiffness within spheroids, micropipette aspiration was used and indicated a mostly reversible stiffness of *E*_sph_ ∼ 0.7 kPa (**Figure 1C**). This is similar to the stiffness of a healthy and highly cellularized soft tissue such as brain tissue that lacks ECM ^2^.

**Figure 1.**
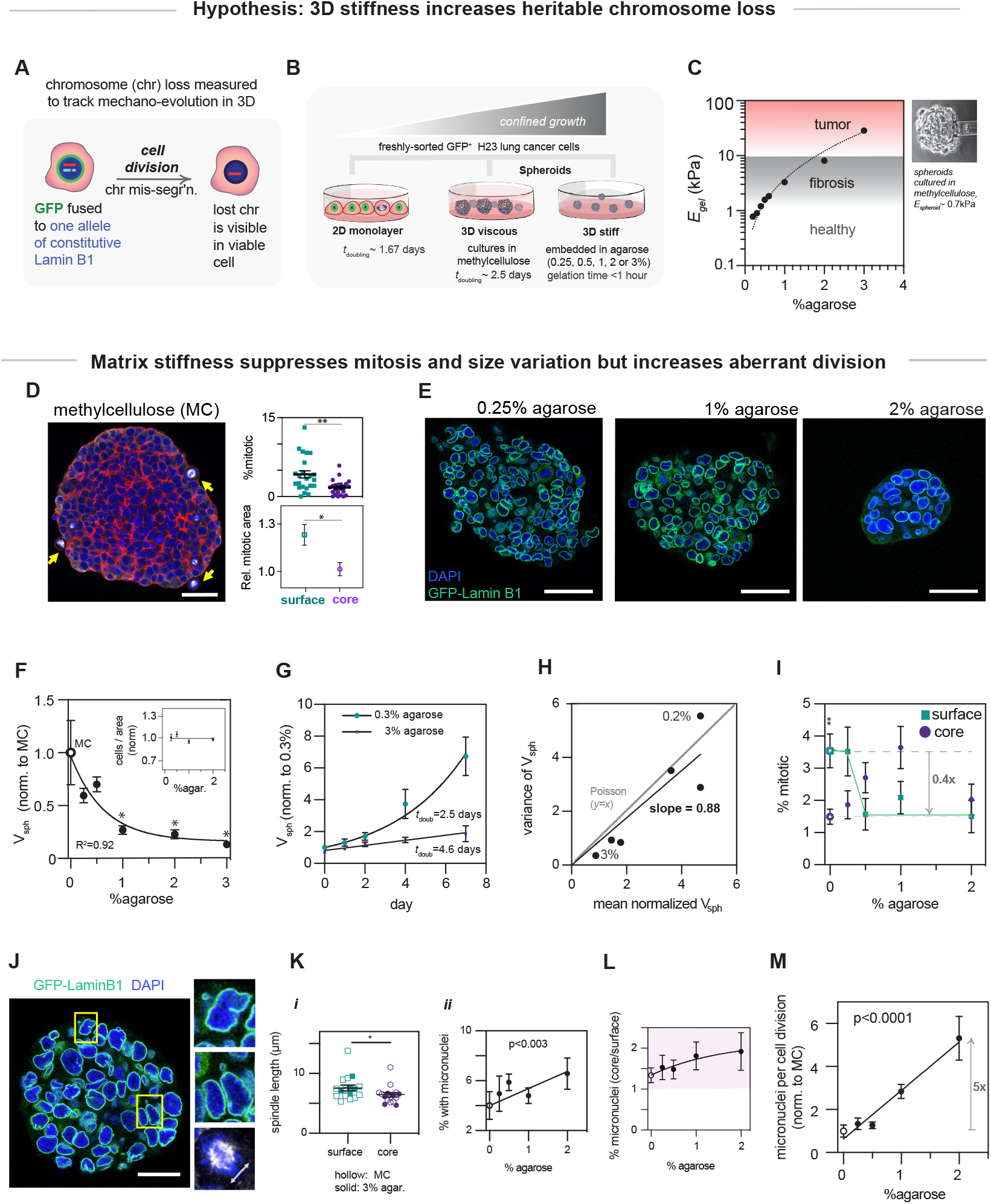
Matrix stiffness suppresses mitosis but increases aberrations per cell division. **(A)** Schematic of the chromosome reporter (ChReporter) cell line demonstrating GFP fused to one allele of the constitutive Lamin B1 gene on Chromosome 5 of human H23 lung cancer cells. **(B)** Freshly sorted GFP-positive H23 cells cultured as 2D monolayers or as spheroids in viscous methylcellulose or in agarose hydrogels of varying stiffnesses. **(C)** Elastic moduli **(E)** of agarose hydrogels that correspond to lung tissue that is healthy (0.2% and 0.3%), fibrotic (0.5% and 1%), or tumor-like (2% and 3%) ^33^. The stiffness of an H23 spheroids cultured in methylcellulose measured using micropipette aspiration (top right). **(D)** Confocal fluorescence cross-section of an H23 spheroid cultured in methylcellulose, showing more mitotic cells on the surface than in the core. Red, F-actin; white, α/β tubulin; blue, DNA. Scale bars, 50 μm. Mean ±SEM for 3 independent experiments. **p<0.01 (t-test). **(E)** Confocal image cross-sections show growth is suppressed for H23 spheroids in stiff agarose hydrogels for the same culture times. Scale bars, 50 μm. **(F)** Quantification of spheroid volume for spheroids cultured in methylcellulose (MC) or agarose hydrogels of varying stiffnesses. Inset: Cell density is unaffected by agarose density based on quantification of cells per cross-sectional area normalized to MC. Shown are mean ±SEM for 3 independent experiments. *p<0.05 (one-way ANOVA). **(G)** Growth curves of spheroids cultured in soft (0.3% agarose) or stiff (3% agarose) gels show that stiffness suppresses growth. Shown are mean ±SEM for 3 independent experiments. **(H)** Spheroid-to-spheroid size variation is lowest in stiff hydrogels (based on panel-F). **(I)** In soft gels, mitotic cells are more common on the surface than in the core, but they are equally low in stiff hydrogels. Shown are mean ±SEM for 3 independent experiments. **p<0.01 (two-way ANOVA). **(J)** Immunofluorescence image of a spheroid cultured in methylcellulose demonstrating micronuclei and a metaphase spindle. Scale bars, 50 μm. **(K) *i***. The metaphase spindle of cells near the core of a spheroid is shorter compared to the cells on the surface. Hollow symbols, methylcellulose; solid symbols, 3% agarose. *p<0.05 (t-test). ***ii***. Micronuclei in spheroids tend to increase from methylcellulose to stiff agarose gels. p<0.003 for the liner fit. Shown for both are mean ±SEM for 3 independent experiments. **(L)** The percentage of cells with micronuclei is higher at the core of a spheroid. Shown are mean ±SEM for 3 independent experiments. **(M)** Quantification of micronuclei normalized with respect to spheroid volume to account for differences in spheroid growth in different hydrogels. Shown are mean ±SEM for 3 independent experiments. (p<0.0001 for the linear fit)

To assess the expected suppression of growth found for other cell types in spheroids surrounded by a high 3D stiffness ^14^, H23 spheroids were grown in various %agarose gels with stiffness up to ∼40-fold higher than that measured for the spheroids themselves. At high agarose concentrations (1-3%) that mimic the stiffness of fibrotic and tumor tissues, the gels are more elastic than viscous, and spheroids were smaller and slower growing than in methylcellulose (**Figure 1E-G**). This confinement result is intuitive. At low agarose densities (0.2-0.3%) near the gelation point where the agarose was a viscous fluid during handling, spheroids were about the same size as those cultured in methylcellulose. Nonetheless, mean cell density in spheroid cores proved independent of agarose density (**Figure 1F, inset**).

What has not been previously emphasized in spheroid studies and is important to findings below for chromosome loss, is that the stiffest gels minimize the *variance* in spheroid size (**Figure 1H**). Moreover, across different agarose concentrations, size variation was proportional to mean size, consistent with a Poisson process of rare events such as cell division, rather than a more random Gaussian process. Mitosis in standard 2D cultures of both normal and tumor-derived lung cells has been shown to follow Poisson statistics ^34^. We indeed find that as matrix stiffness increases, mitosis shifts from being mostly at the surface (where 3-4% of cells are mitotic) to being uniformly low throughout the core (**Figure 1I**). The surface-to-core mitotic ratio is ≤1 and shows the highest variance at hydrogel stiffnesses > 1 kPa. This observation aligns with our measurement of *E*_sph_ ∼ 0.7 kPa and again suggests that higher stiffness suppresses mitosis on the spheroid surface. Increased mitosis in the core from 0% to 1% agarose seems analogous to increased proliferation on stiff 2D substrates including plastic (e.g. **Figure 1B**) but might also reflect prolonged division under confinement ^35^. However, in 2-3% agarose, mitosis is more clearly and uniformly suppressed by the more rigid 3D surroundings.

Despite this suppression of mitosis in stiff hydrogels, these spheroids generally display surprisingly high levels of micronuclei (**Figure 1J-M)**. Micronuclei typically result from chromosome mis-segregation during division ^8,9^, and for 2D cultures, external compression of the mitotic spindle can drive viable mis-segregation ^16^. We therefore measured mitotic spindle lengths and discovered core spindles are shorter than surface spindles (**Figure 1K-I**: by 10-15% ≈ 0.87-fold), consistent with smaller cross-sectional areas for core divisions (**Figure 1D**: (0.8-fold)^1/2^ = 0.89-fold). Stiff agarose showed the same surface versus core difference in spindle length despite the suppression of surface divisions. Importantly, micronuclei also tend to be more prevalent in the spheroid core than the surface – regardless of stiffness (**Figure 1L**). Micronuclei can accumulate DNA damage and trigger cell cycle checkpoints ^36^, which could explain some growth suppression observed in the spheroid cores and stiff hydrogels.

To account for differences in rates of division that sometimes generate micronuclei (**Figure 1F-H**), we normalized the percentage of micronuclei to total divisions (i.e. spheroid size) and find a 5-6 fold increase in micronuclei frequency per division in the stiffest gels (**Figure 1M**). Micronuclei in one daughter cell indicates chromosome loss in the other daughter cell ^8,9,37^, which suggests that spheroids in stiff gels will exhibit more frequent chromosome loss. In addition, despite the slow growth and low variance of spheroid size in stiff gels (**Figure 1F-H**), a high *variance* in micronuclei for spheroids and spheroid cores for stiff gels (**Figure 1L**,**M**), also suggests a high variance in chromosome loss between spheroids.

### Matrix stiffness increases ChReporter loss per division & enhances inter-spheroid variation

To investigate chromosome loss, fluorescence-activated cell sorting for ChReporter-positive cells was done before each experiment. Loss of GFP signal at the nuclear envelope in confocal microscopy sections was evident in individual cells and colonies that suggested heritable loss (**Figure 2A**). These ChReporter-negative cells were isolated using fluorescence-based sorting and used to confirm chromosome loss by genetic analyses (not shown). This rules out epigenetic processes associated with microenvironmental factors such as matrix stiffness that are reversible over many days ^2^.

**Figure 2.**
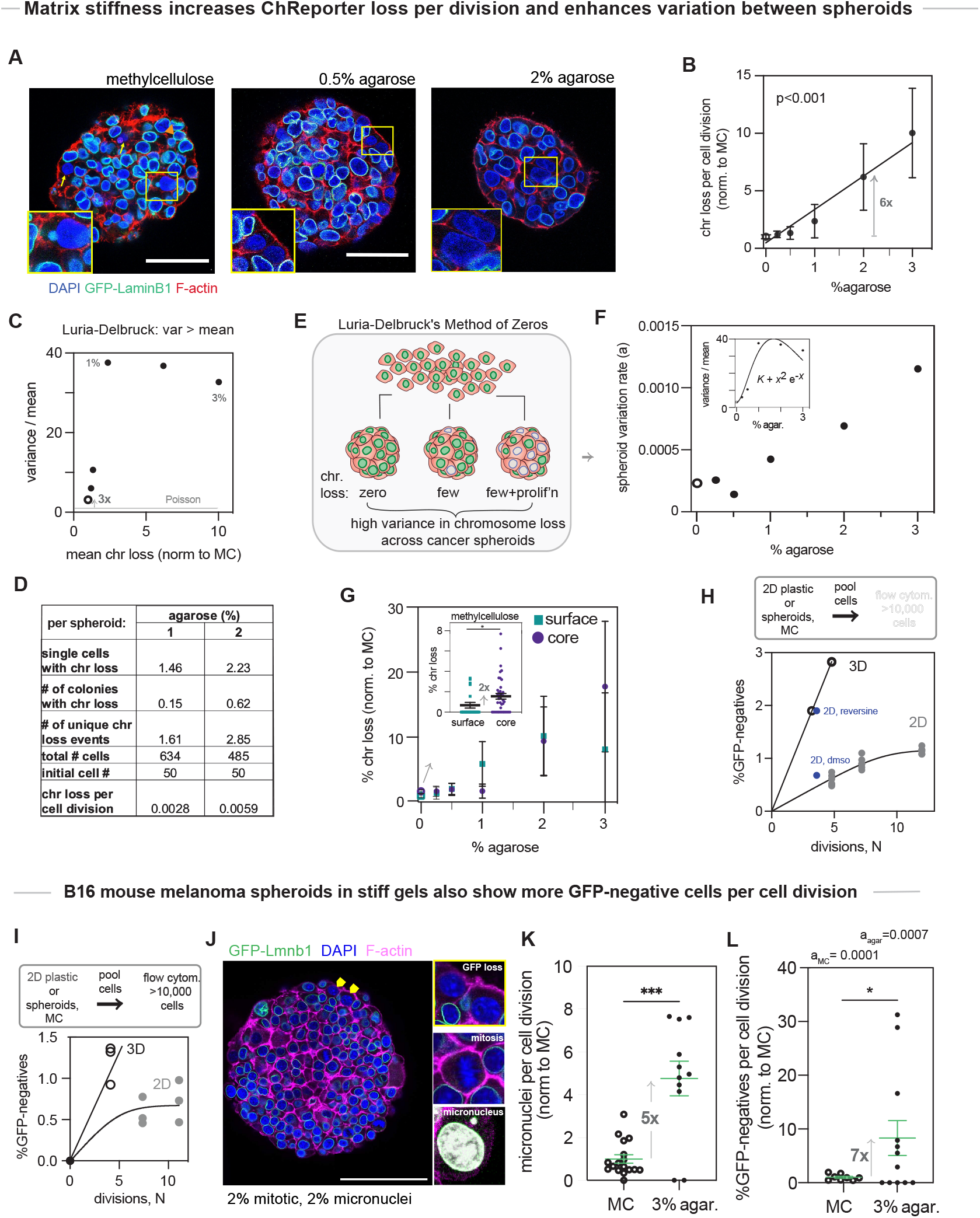
Matrix stiffness increases ChReporter loss per division and enhances chromosome variation between spheroids. **(A)** Fluorescence images of spheroids cultured in hydrogels with different stiffnesses demonstrate that matrix stiffness increases chromosome loss rate. Scale bars, 50 μm. **(B)** Chromosome loss normalized with respect to spheroid volume to account for differences in cell division. Shown are mean ±SEM for 3 independent experiments. (p<0.001 for linear fit) **(C)** Variance / mean of chromosome loss always exceeds unity to reveal a non-Poisson process modulated by stiffness. **(D)** The frequency of chromosome loss per cell division for spheroids cultured in 1% versus 2% agarose hydrogels, demonstrating that stiffness increases the frequency of chromosome loss. (**E**,**F**) The method of zeros proposed by the Luria and Delbruck model allows for a separate measurement of variation rate, with the plot showing that stiff hydrogels increase chromosome loss. **Inset:** The Luria and Delbruck model estimates the (Variance/Mean) as the product of the variation rate, *a*, and divisions, N, and *a* is approximately a quadratic function of % agarose whereas N decays exponentially with % agarose per Figure 1F. **(G)** Chromosome loss increases for cells near the core of a spheroid cultured in methylcellulose compared to those on the surface (±SEM for 3 independent experiments; *p<0.05 with t-test), whereas this trend disappears for spheroids cultured in agarose hydrogels. (**H, I**) Flow cytometry analysis of H23 cells or mouse B16 melanoma cells cultured on a 2D plastic surface or as spheroids in MC reveals that the 3D microenvironment increases the chromosome loss rate (Welch**’**s t-test, ****p<0.0001). Addition of reversine to H23 2D cultures increases chromosome loss rate in 2D similar to H23 Spheroids, based on mean ±SEM for 3 independent experiments. (**J**) Immunofluorescence staining of B16 mouse melanoma spheroids demonstrating mitotic cells, GFP loss including a GFP-negative colony, and micronucleus formation. (**K**,**L**) Micronuclei per cell division increase in B16 spheroids cultured in stiff agarose, as do GFP-negative cells and variation rate *a*, based on mean ±SEM for 3 independent experiments.

A ∼6-fold increase in chromosome loss per division is found for spheroids within stiff 2% agarose gels relative to viscous methylcellulose (**Figure 2B**). This aligns well with the increase in micronuclei per division (**Figure 1M**). A proportionality between ChReporter loss and micronuclei was also demonstrated in 2D cultures of both H23 cells and A549 lung cancer cells treated with reversine (not shown), which is an MPS1i drug that induces chromosome mis-segregation ^21,29^. In the stiffest gels of 3% agarose, the percentage of ChReporter-negative cells was even higher than in 2% agarose gels, despite imaging at a timepoint (day 7) when the starting cell population (∼50 cells) had undergone only about one division. The softest gels and methylcellulose (**Figure 1G**) allow for more frequent cell divisions that increase the likelihood of errors, which underlines the importance of normalizing to the total number of cell divisions.

Variance in chromosome loss per division across different spheroids was considerably higher (∼100-fold) in the two stiffest gel conditions compared to methylcellulose and the softest gels. Importantly, the ratio (variance/mean) for all conditions was greater than 1 (with 1 expected for Poisson statistics, **Figure 2C**), which is consistent with Luria-Delbruck**’**s seminal theory on mutational variation applied to their parallel cultures of bacteria ^20^. A high variance results from non-random proliferation of mutants (i.e. colonies) after a rare and *irreversible* genetic event that does follow Poisson statistics. We find that even in methylcellulose which shows a minimum ratio for variance/mean of ∼3, heritable chromosome loss in GFP-negative colonies supports the theory (**Figure 2A**). The initial increase in this ratio with agarose (up to a 6-fold increase in 2% agarose) is followed by a slight decrease that is consistent with fewer divisions in the stiffest gels (**Figure 1F**). No divisions would give no genetic change.

For an absolute quantitation of chromosome loss rate per division, we focused on the spheroids cultured in the stiffest gels where assessments of GFP-negative cells were made after only 1-2 doubling times compared to the many doubling times in methylcellulose (**Figure 1G**). Longer incubation times allow for more proliferation of GFP-negative cells, potentially overestimating chromosome loss rates per division. Importantly, we counted each colony as a single GFP-negative event when calculating the number of unique loss events. By converting spheroid volume to cell number and again estimating total divisions (N_t_), we determined a frequency of 0.59% loss of the ChReporter allele per division in 2% agarose, and about half this rate in 1% agarose (**Figure 2D**). If this same rate applies to all ∼60 chromosomes in H23 cells, then heritable chromosome loss in stiff gels occurs in roughly one-third of all divisions, which is high but not ubiquitous.

Importantly, across all conditions, a fraction of the imaged spheroids did not display any GFP-negative cells (**Figure 2E**), which is consistent with the fact that chromosome loss is inherently a rare event. According to Luria-Delbruck**’**s **‘**Method of Zeros**’**, the frequency of independent cultures (or spheroids) without any genetic changes can be used as a second way to estimate mutation rates. The lower bound of the variation rate is *a* = - ln *P*_0_ / N_t_, where *P*_0_ is simply the fraction of spheroids showing no chromosome loss ^20^. The calculated rate was plotted for different agarose concentrations and methylcellulose (**Figure 2F**), and the increase with stiffness aligns with the previous trend obtained from counting individual GFP-negative cells (**Figure 2B**). Differences in absolute values between methods are expected according to Luria-Delbruck with *a*_*l*_ being an underestimate ^20^. Nonetheless, the variation rate for stiff gels (∼0.1% per division) is within an order of magnitude (**Figure 2D**). Based on Luria-Delbruck**’**s theoretical framework (see Methods), a quadratic fit of variation rate *a* was applied, combined with decreased cell divisions in stiffer gels (**Figure 1F**), to explain the suppressive effect of high matrix stiffness on ChReporter loss in terms of (variance/mean) (**Figure 2F-inset**).

Imaging in this study is at the sub-organelle level, providing high resolution measurements of genetic change in 3D with single-cell sensitivity at better than 1-per-1000 cells, which is more sensitive than most sequencing methods. This allowed us to observe, for example, that the percentage of GFP-negative cells was approximately two-fold *higher* in the more confined core of spheroids cultured in methylcellulose compared to losses on the surface (**Figure 2G-inset**). This aligns with increased spindle compression and more micronuclei in the core (**Figure 1K**,**L**). The lack of a clear trend in surface-versus-core chromosome loss across the different agarose gels (**Figure 2G**) could reflect the increasing variance in core-to-surface ratio of micronuclei, which suggests that surface cells experience confinement similar to core cells in stiffer gels – consistent with the equally suppressed mitosis in the stiffest gels (**Figure 1I**).

To confirm the imaging-based results and to increase the throughput (at the expense of seeing colonies), we used flow cytometry to analyze >10,000 cells from standard cultures and from spheroids in methylcellulose cultures (**Figure 2H**). Flow cytometry showed ∼4-5 fold more GFP-negative cells than 2D cultures after an equal number of N=5 divisions (N_t_ = 2^N^). Loss is monotonic with division in both cases but increases to a greater extent in spheroids and saturates in 2D at a level of ∼1% that is similar to that for cells imaged on the surface of spheroids cultured in methylcellulose (**Figure 2G-inset**). Flow cytometry is a bulk assay that cannot distinguish cells in the surface layer from cells in the core. However, because there are more core cells in large spheroids such as in methylcellulose (see Methods: *Surface layer and Core cell numbers*), a high GFP-negative result from flow cytometry on spheroids reflects the higher loss rate in the core, which is consistent with image analysis.

Reversine treated 2D cultures show increased GFP-negative cells to a similar extent as 3D (**Figure 2H**), consistent with micronuclei induction. Interestingly, reversine added to 3D spheroids shows no effect (not shown). The effect of such MPS1i**’**s relies on an intact microtubule spindle to drive mis-segregation. Hence, the lack of effect in 3D seems consistent with a strong perturbation of the microtubule spindle within the spheroids (**Figure 1K**) that thereby limits the effect of reversine.

Applying Luria-Delbruck models to the flow results cannot account for colony formation (see Methods and fits to (**Figure 2H**) but gives *a*_*u*_ = 0.0081 changes per division for 3D (MC) versus *a*_*u*_ = 0.0023 for 2D. These are overestimates that are at least in the range of rates determined from image analyses of H23 spheroids (**Figure 2D**), and the 4-fold increase from 2D to 3D seems in line with effects of confinement within a spheroid.

### Spheroid results with different cells, species, chromosome

To assess ChReporter loss in another solid tumor cell type as well as another chromosome, we chose mouse melanoma B16 cells and the same GFP-tagged gene (*Lmnb1*), which happens to be on the much smaller chromosome, Chr18 (see Methods). We employed flow cytometry for our first assays on parallel cultures in 2D and spheroids in methylcellulose, and the results showed ∼2-3 fold more GFP-negative cells than cells grown on rigid tissue culture plastic that exhibited ∼0.5 - 1% GFP-negative cells (**Figure 2I**). The result is similar to H23 results (**Figure 2H**), and reversine in 2D cultures again increased GFP-negative cells to an extent that phenocopied the results for 3D spheroids (not shown). Applying Luria-Delbruck models to the flow results gives for 3D (MC) *a*_*u*_ = 0.0042 changes per division compared to *a*_*u*_ = 0.0016 for 2D, which are only slightly lower than the rates also determined by flow for H23 spheroids (**Figure 2H**).

Imaging of the B16 spheroids cultured in methylcellulose revealed colonies of GFP-negative cells along with the presence of mitotic cells and micronuclei (**Figure 2J**). The levels of mitotic cells and micronuclei were slightly lower than those observed for H23 spheroids (**Figure 1I**,**K**), consistent with the slightly lower *a*_*u*_. In stiff agarose gels, B16 spheroids showed about half as many mitotic cells compared to methylcellulose (not shown), while micronuclei counts per cell division were 5-fold higher (**Figure 2K**), consistent with H23 results (**Figure 1M**).

Importantly, ChReporter loss per division increased by ∼7-fold, as indicated by both GFP-negative cell counts and the **‘**Method of Zeros**’** variation rate (*a*) (**Figure 2L**). The latter is in reasonable agreement with estimates of *a*_*u*_ from flow (**Figure 2I**).

Cell lysates and genomic DNA from the sorted and expanded GFP-negative B16 population were analyzed by immunoblots and PCR, which provided more molecular evidence for loss of the GFP-tagged allele (not shown). To investigate whether GFP-negative B16 cells also emerge in 3D solid tumors in vivo, freshly-sorted GFP-positive B16 cells were implanted subcutaneously in mice, and tumors were harvested after 19 days for flow cytometry analyses. The GFP-negative B16 cells increased from a steady state 0.29% in long-term 2D culture by almost 2-fold in the large 3D masses (not shown), which is modestly lower than in vitro findings for B16 cells (**Figure 2I**). Genomic DNA from the GFP-negative B16 cells generated in vivo again provided molecular evidence for loss of the GFP-tagged allele. These results thus align with the extensive results using the human H23 cells.

### Matrix stiffness increases colonies with ChReporter loss

Cell death or quiescence can be triggered by chromosome mis-segregation, but a viable and heritable genetic change is indicated by colony formation (**Figure 3A**). Information such as this for initially rare genetic changes is extremely challenging for sequencing methods and standard karyotyping (on 50-100 cells, for example). We find for stiffer gels that 60% of H23 spheroids exhibited one or more colonies with ChReporter loss, while spheroids in softer gels, including methylcellulose, most frequently showed no colonies or just one colony (**Figure 3B**). These colony formation results align with the larger variation in spheroids in stiffer matrices relative to softer matrices noted in the earlier analyses of GFP-negative counts (**Figure 2B**,**C**), and this is particularly important but surprising because it occurs despite suppressed growth in stiffer matrices (**Figure 1E-I**). Nonetheless, for stiff gels, a very early chromosome loss would allow for ∼2 doublings (to 4 cells) but not 3 doublings (8 cells), which is consistent with the observed maximum of 5-6 cells (3 colonies in **Figure 3C**).

**Figure 3.**
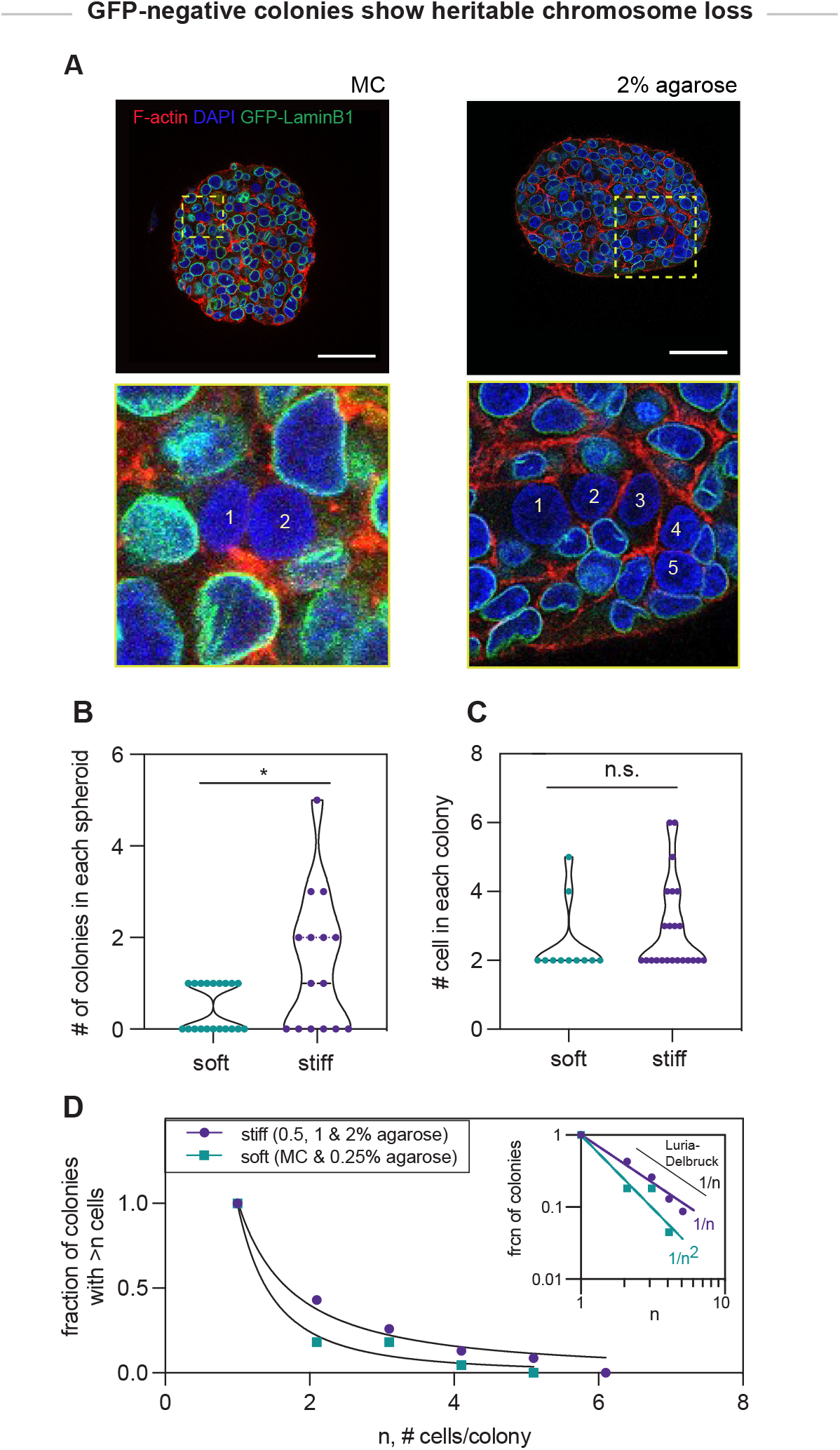
Colonies of GFP-negative cells reveal heritable chromosome loss. **(A)** H23 spheroids cultured in methylcellulose or agarose stained for F-actin and DNA display colonies of GFP-negative cells, which reveal that **(B)** the number of colonies and **(C)** the number of GFP-negative cells in each colony increases in stiff matrices. Shown are mean ±SEM for 3 independent experiments. *p<0.05 (Kolmogorov-Smirnov test). **(D)** Spheroids cultured in stiff agarose hydrogels harbor a higher fraction of larger GFP-negative colonies.

Colony sizes also tended to be larger in stiffer matrix conditions (**Figure 3C, D**), which is most evident as a ∼1/n decay of colony counts versus colony size (**Figure 3D - inset**). Such a decay is consistent with recent studies of mutating *E*.*Coli* where colonies of ∼100 mutant cells or more also fit this ∼1/n expectation from Luria-Delbruck^21^. Overall, the colony counts are consistent with increased ChReporter loss in stiff gel conditions and demonstrate that this genetic change can be propagated over generations, which again aligns with our application of Luria-Delbruck**’**s basic models of proliferation after a rare genetic induction event ^20^.

### Suppressing Myosin-II induces chromosome loss in 3D Spheroids but not in 2D

Tumor suppressors impact growth moreso in 3D than standard 2D cultures and include the molecular motor myosin-II ^23^. Myosin-II is ubiquitous and well known to sense matrix stiffness ^2^. We hypothesized that suppression of such a 3D-specific tumor suppressor can increase chromosome loss rate even without affecting growth. In 2D cultures, the myosin-II inhibitor blebbistatin indeed shows no effect on a low basal rate of chromosome loss (not shown), even though knockdown or deletion of myosin-IIA (MYH9) *initiates* tumors in mouse skin ^26^. Myosin-II suppresses the PI3K-AKT oncogene pathway ^38^ that also involves PTEN which suppresses aneuploidy ^32^.

Blebbistatin treatment of spheroids in methylcellulose and stiff agarose gels tended to *suppress* spheroid size even though differences were not statistically significant (**Figure 4A, B**) and neither were mitotic counts (not shown). However, staining for F-actin showed some cells lacked any cortex between adjacent cells, indicating multinucleation, and blebbistatin led to ∼8% multinucleated cells compared to <1% with DMSO for spheroids in methylcellulose. For stiff gels, blebbistatin and DMSO both resulted in ∼3-4% multinucleation (**Figure 4C**). Myosin-II deletion in suspension cultures of amoeboid cells causes multinucleation through cytokinesis failure, which is a known pathway to chromosomal instability and losses ^11,32,39^.

**Figure 4.**
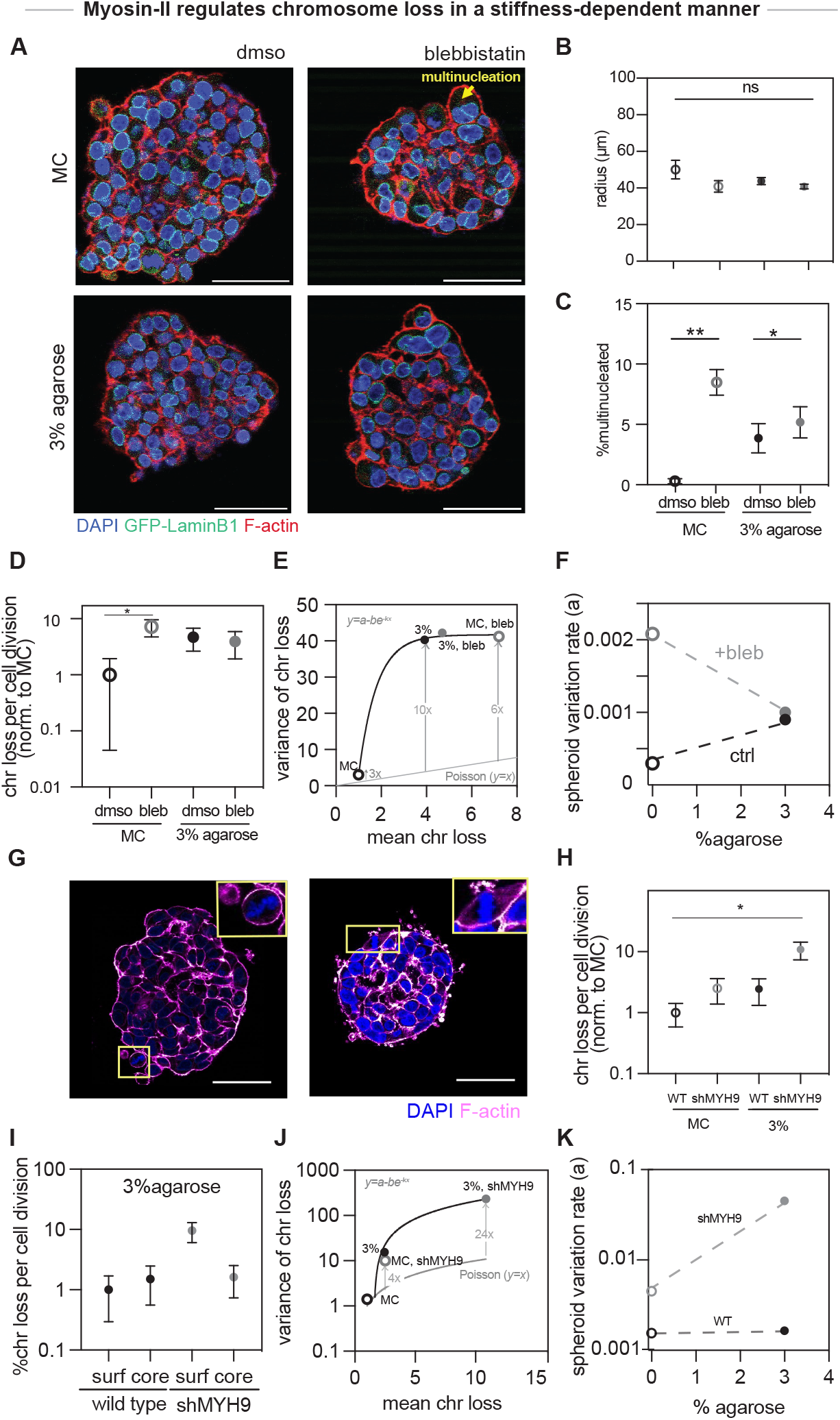
Myosin-II regulates chromosome loss in a stiffness-dependent manner. **(A)** Fluorescence images of H23 spheroids treated with blebbistatin. (**B**,**C**) Blebbistatin treatment does not affect the size of spheroids, but increases multinucleation in spheroids cultured in methylcellulose. Shown are mean ±SEM for 3 independent experiments. *p<0.05, **p<0.01 (Welch**’**s ANOVA test using Dunnett**’**s test). (**D-F**) Inhibiting Myosin-II with blebbistatin increases chromosome loss in spheroids cultured in methylcellulose. Shown are mean ±SEM for 3 independent experiments. Variance versus mean of chromosome loss is greater than one, which reveals a non-Poisson distribution and demonstrates that variance increases with stiffness and blebbistatin treatment. The method of zeros from Luria and Delbruck also shows blebbistatin treatment increases chromosome loss. (**G-K**) Immunofluorescence images of control or shMYH9 H23 spheroids cultured in stiff agarose hydrogels. Knockdown of MYH9 increases chromosome loss per cell division in all spheroids, and Chromosome loss increases in shMYH9 cells on the surface of spheroids cultured in stiff agarose hydrogels. Shown are mean ±SEM for 3 independent experiments. *p<0.05 Variance versus mean chromosome loss exceeds one, which reveals a non-Poisson distribution and demonstrates that stiffness and MYH9 knockdown increase variance. Spheroid variation rate increases in shMYH9 spheroids cultured in stiff agarose hydrogels. Scale bars, 50 μm.

Chromosome loss was greatest in blebbistatin-treated spheroids in methylcellulose and intermediate in stiff 3% agarose gels, independent of blebbistatin (**Figure 4D**). The variance in chromosome loss between spheroids was again ∼3-fold above Poisson statistics for control spheroids in methylcellulose and increased to 6-10 fold above Poisson for blebbistatin-treated and/or stiff gel conditions (**Figure 4E**). This trend aligns with random chromosome loss followed by proliferation. The GFP-negative results indicate a ∼7-fold higher chromosome loss rate with blebbistatin in methylcellulose, consistent with the variation rate *a* obtained using Luria-Delbruck**’**s **‘**Method of Zeros**’** from the fraction of GFP-positive spheroids (**Figure 4F**). Our finding that myosin-II inhibition with blebbistatin has no effect on the low basal level of chromosome loss in 2D cultures (not shown) thus highlights a strictly 3D effect of this candidate tumor suppressor on chromosome instability.

MYH9 or nonmuscle myosin-IIA is the widely expressed myosin-II proposed to be a tumor suppressor for skin cancer based on knockdown in mice ^26^, but mechanisms have remained understudied. We applied a similar shMYH9 knockdown strategy in human H23 cells, noting that these cells express multiple myosin-IIs that can all be inhibited by blebbistatin (not shown). Knockdown resulted in flattening of mitotic cells on the spheroid surface within stiff gels (**Figure 4G**), consistent with myosin-II contributing a major fraction (∼70% or more) of the cortical tension that drives mitotic rounding ^39^; this is important because chromosome mis-segregation can be driven by external compression of the mitotic spindle in 2D culture ^16,35^.

Indeed, a 5-10 fold increase in GFP-negative cells per division in stiff gels is evident mostly on the spheroid surface (**Figure 4H, I**). The variance in GFP-negative cells once again exceeded controls and Poisson statistics for knockdown cells (**Figure 4J**), consistent with a higher rate of chromosome loss, and given that overall growth differences were not significant (not shown). Finally, Luria-Delbrück**’**s **‘**Method of Zeros**’** analysis of the fraction of purely GFP-positive spheroids highlighted a greater than10-fold increase in variation rate *a* in stiff gels following the knockdown of MYH9 (**Figure 4K**).

## Discussion

### Chromosome loss has mechanobiological determinants

To maximize successes for immune therapies or other treatments, even rare mutations should be better understood in order to better anticipate their effects and future recurrence. Mutations induced here by mechanobiology factors of ECM stiffness and contractility appear novel even though an optimal stiffness for cytoskeletal organization and gene expression is already clear from diverse studies of stem cells in 2D ^2^ and cancer cells in 3D ^1,40^. Our main finding that ECM stiffness drives a higher rate of chromosomal copy number changes in 3D cancer spheroids – for standard human and mouse cancer lines – not only explains broad trends from liquid to solid tumor types in humans (**Table 1**) but also the high heterogeneity of solid tumors. Our findings for the variance as non-Poisson and non-Gaussian (**Figures 2C,F** and **4E**,**J**) indeed indicate induction and proliferation of mutants that are foundational for subsequent selection in mechano-evolution. Mechanics can indeed modulate mutation rates, but rate effects are similar for very different chromosomes rather than specific (for determinism of Lamarck), and so mechano-evolution seems more of a *combined* Darwin-Lamarck process.

Our finding that myosin-II disruption further drives chromosome loss (**Figure 4**) underscores a mechanobiological contribution even though other tumor driver genes in the same pathway are selected for more frequently in human cancers ^38^. Myosin-II knockout in yeast does lead to aneuploidy and possibly through selection of rare mutants ^41^; knockdown in relatively stiff mouse skin leads to cancer ^26,27^. Skin is a moderately stiff tissue with abundant ECM ^4,42^, and so a context dependence in our results for myosin-II as a suppressor of tumor evolution is reasonable. Myosin-II also has an essential role in cell division for liquid suspension cultures but not standard 2D substrates where migration facilitates division ^25,43^. Our H23 results *conceptually* align with screening results for 3D-specific cancer driver genes ^23^, but the latter results indicated spheroids should grow faster with myosin-II disruption – which we do not observe (**Figure 2E**). Indeed, chromosome loss occurs in the same or fewer divisions *N* and is best explained by perturbed mitosis in 3D. Similar studies of tumor suppressors beyond myosin-II as well as oncogenes in 3D may yield similarly surprising results – with a key signature likely being the ongoing induction of heritable chromosome number changes as small colonies.

### Human patient: Intra-tumor variance of diverse Chromosome Losses

A spatially-resolved snapshot of one patient**’**s solid tumor (Liver Hepatocellular Carcinoma, LIHC) was obtained by sequencing 23 sites of ∼20,000 cells per site in a section of ∼35-mm diameter (not shown) ^48^. Losses exceed gains in their table of CNVs, and *all* sites showed loss of most of Chr-5, 13, and 16 which indicates a strong Darwinian selection after an ancestral loss.

However, some chromosome losses (e.g. Chr**’**s 7 and 19) occur only as small colonies, which supports the authors**’** conclusion that extremely high and ongoing mutational diversity points to non-Darwinian cell evolution in the tumor ^48^. Sub-clonal differences suggest application of the Method of Zeros (**Figure 2E**) for each chromosome at all 23 sites, leading to a chromosome-average mutation rate *a* ∼ 10^−5^ per division; this is ∼10-fold lower than our results for human H23 and mouse B16 cells in methylcellulose and soft gels (**Figure 2**). The human liver tumor also seems representative in showing losses or gains for 23% of the genome (3.2 Gb normally), which is within 1 S.D. of the mean CNV fraction for all LIHC patients in **Table 1**. Broader trends for tumor variation across patients were therefore assessed.

### Inter-tumor variance associates with mean copy number changes and high collagen

To further relate to human cancers our findings that the highest chromosome loss per division and the highest variation occur in the stiffest 3D microenvironments, we further analyzed public data for chromosomal aberrations quantified in The Cancer Genome Atlas (TCGA) (per **Table 1**) and for purified human tumor lines in the Cancer Cell Line Encyclopedia (CCLE). Black on white heatmaps help illustrate our approach by displaying deviation of chromosome numbers or segments thereof for individual patients with four specific cancer types (**Figure 5A**). Patients with leukemia (LAML, a liquid cancer) show just a sparse set of chromosomal changes, whereas ovarian cancer patients (OV) have solid tumors with high copy number variation and lung and pancreatic cancer patients (LUAD, PAAD) fall in between. The copy number variations (CNVs) must be heritable within the patient**’**s tumor given the dominance of the signal, and for a given type of tumor, the mean fraction of the genome with CNVs correlates with collagen-I expression as a median of *COL1A1* and *COL1A2* (**Figure 5B, Table 1**).

**Figure 5.**
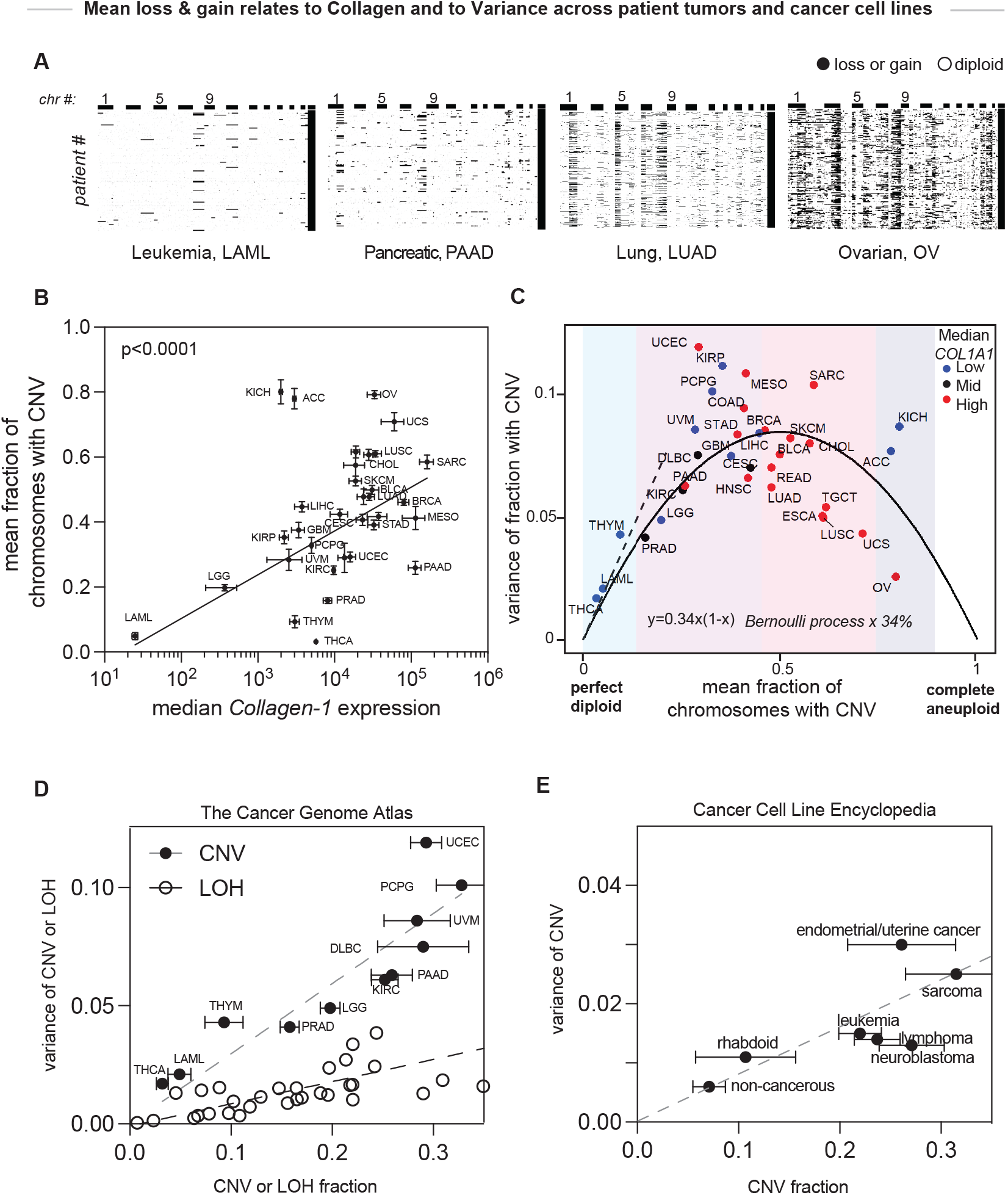
Chromosome losses and gains across patient tumors increase with collagen-I, and the variance also tracks with the mean. **(A)** Chromosome gains and losses in acute myeloid leukemia (LAML), pancreatic adenocarcinoma (PAAD), lung adenocarcinoma (LUAD) and ovarian cancer (OV) from the Cancer Genome Atlas (TCGA). (**B**,**C**) Fraction of genome displaying chromosome number variation (CNV) and variance in CNV plotted with median Collagen-I expression. Data is from TCGA. (**E**,**F**) Linear range of variance in CNV and variance in Loss of Heterozygosity (LOH) plotted versus the respective means, and for data from TCGA and purified cultured cells of the Cancer Cell Line Encyclopedia (CCLE).

Importantly, patient-to-patient differences lead to a variance between tumors of a given type, which is not random per Gaussian statistics. Instead, the variance increases with CNV until about half the genome shows alterations and then the genome becomes saturated by CNV (**Figure 5C**,**D**). Low collagen tumors including well-known liquid tumors (e.g. leukemia, LAML) exhibit low variance as well as low CNV. Remarkably, the pan-cancer trend for inter-patient variance is modeled well by a Bernoulli distribution for each cancer type but with inclusion of a common prefactor:

Variance = α Mean * (1 – Mean) with α_TCGA_ = 34% (**Figure 5C**)

A Bernoulli distribution has α = 100%, as does a Poisson distribution that applies for Mean <<1. However, the lower variance of 34% suggests a surprisingly constant fraction of down-selection from standard statistics to heritable CNV across all cancer types.

A similar trend for the variance between human tumor lines of a given cancer type in the CCLE is also evident in the low CNV range that includes non-cancerous cells and leukemia (**Figure 5E**). A smaller α ∼ 10% suggests more homogeneity in these standardized 2D cultures compared to intact 3D tumors. Nonetheless, prolonged cell culture is well-known to impact aneuploidy, with many solid tumor lines becoming approximately triploid ^44^. While loss of heterozygosity (LOH) measures a sequence specific loss, it conforms to LOH ∼ ½ CNVs ^45^ and exhibits s variance across cancers that also scales linearly with its mean (**Figure 5D**). Patient survival based on LOH and CNV might also correlate with the variance for a given a tumor type, although further study is needed.

Despite a major role for selection in the evolution of the tumors curated in TCGA and CCLE, the variance in chromosome number differences between established human tumors of a given type generally relates to the mean – *and* generally increases with ECM stiffness according to our experiments which fit well to Luria-Delbruck type statistics for initial genetic change (**Figures 2-4**). This includes the 1/n decay in the frequency of colonies with n cells in stiff gels (**Figure 3D**) that was postulated to apply to solid tumors based on theory for much larger colonies of mutating *E*.*Coli* plus some key assumptions such as no cell death, no migration, no selection, and uniform growth ^20,21,46^. In our system, stiff ECM constrains mitosis (uniformly at the surface and in the core per **Figure 1I**), and B16 spheroids show minimal migration in a few initial movies – although further study is needed to possibly explain the 1/n^2^ decay in soft gel conditions. Selective advantages should apply to chromosome loss, but our recent ChReporter study that ties faster growth to loss of a tumor suppressor ^44^ shows the effect is small over just a few divisions as studied here. Many other *in vivo* factors including immunity no doubt influence the inter- and intra-tumor variation in such genetics, with implications for treatment – but even immune responses are sensitive to ECM ^47^.

### Limitations of the study

Genetic changes help define tumor heterogeneity that challenges many treatments, but mitotic mistakes often cause cell arrest or death that limits heterogeneity. The microscopy approach here illustrates a means of multi-day tracking for rare but visible and viable colonies that relate back to molecular mechanisms of 3D compression of spindles and generation of micronuclei. Our ChReporter-negative cells do reflect loss of protein and thereby mimic what is consequential for loss of a tumor suppressor, but assessing induction of other chromosome losses in additional ECM contexts and more cell lines remains important. A key goal should be generation of larger datasets suitable for statistical extensions of Luria-Delbruck theories to physical mechanisms ^20,21,46^ that relate to human genetics (**Table 1**). Agarose is non-adhesive to cells and enabled a focus on ECM stiffness effects, but studies with collagen-I and other native ECM can help assess cell adhesion effects. An important issue for other ECM**’**s and other cells is whether the spheroids are sufficiently dense and close-packed to cause aberrations in division. Looser aggregates are expected for mesenchymal phenotypes and invasion (e.g. high E-cadherin with low N-cadherin), and these will tend to overcome the needed physical compression that arises from spheroid growth against a surrounding matrix – as already suggested by initial studies of a couple of additional cell lines (not shown). Such issues add to whether apoptosis or cell cycle arrest undermines productive induction of viable, heritable genetic changes.

Lastly, chromosome loss drives the loss of tumor suppressor expression from that particular chromosome which can impact the localization and function of these particularly key proteins in cells (not shown). We also recently demonstrated selection for faster growth using our ChReporter approach ^44^ which underscores the need for further studies of pathways (e.g. p53 pathway) as well as evolved phenotypes. Nonetheless, the short time scales for studies here relative to cell cycle help keep the focus on *induction* of genetic changes in very early heritability rather than *subsequent* selection for dominance of particular clones in established tumors as typically studied (e.g. **Table 1, Figure 5A,B**). A random induction step is the essence of Luria-Delbruck statistics, and yet – across established tumors – what emerges is a cancer-agnostic down-selection from that broad variance (e.g. **Figure 5C-E**).

## Materials and Methods

### Cell culture and reagents

NCI-H23 human lung adenocarcinoma cells were a gift from the laboratory of MC Bassik (Genetics, Stanford University), who reported oncogenic profiles of their H23 spheroids in methylcellulose ^23^, and so we specifically sought their cell line to minimize genetic drift effects. H23 lung adenocarcinoma and B16 melanoma cells (ATCC) were cultured in RPMI medium supplemented with 10% fetal bovine serum (*Sigma*) and 1% penicillin-streptomycin (Gibco). U2OS osteosarcoma and A549 lung adenocarcinoma cells (ATCC) were cultured in DMEM or F12 medium supplemented with 10% fetal bovine serum (*Sigma*) and 1% penicillin-streptomycin (Gibco). For 2D experiments, freshly sorted H23 and A549 cells were cultured in 6-well plates at 50,000 cells per well. MPS1i (Reversine: 0.5 µM for A549 and 1.5 µM for H23; Cayman Chemicals) or blebbistatin (20 µM; Millipore Sigma) were added to fresh culture medium 72 hours after plating. The cells were treated for 24 hours after which the drugs were removed, the cells were washed three times with PBS and fresh media was added. A549 and H23 cells were cultured for an additional 24 and 48 hours respectively, after the drugs were removed. GFP-negative B16 cells were obtained by either sorting for spontaneously generated GFP-negatives after multiple passages or by treating B16 GFP-LaminB1 cells with reversine as previously described.

### ChReporter gene editing

Knock-in chromosome reporter H23 cells were engineered using the CRISPR/Cas9 technology (Roberts et al., 2017). AICSDP-10:LMNB1-mEGFP (Addgene plasmid #87422) donor plasmid construct was used for the GFP-LaminB1 monoallelic chromosome reporter. Using the ribonucleic protein (RNP) technique, recombinant wild-type Streptococcus pyogenes Cas9 (University of California-Berkeley QB3 Macrolab) protein pre-assembled with synthetic CRISPR RNA (crRNA; Horizon Discovery) and a trans-activating crRNA (tracrRNA; Horizon Discovery) duplex were mixed. To introduce donor templates into target cells, we used electroporation (Gene Pulse Xcell Electroporation System, Bio-Rad), where we harvested 700,000 target cells using 0.05% trypsin-EDTA, suspended them in 200 μL of fresh medium without penicillin-streptomycin, and loaded them into a 0.4-cm cuvette. We then added 4 µl each of a 10 µM crRNA:tracrRNA duplex, 10 µM recombinant Cas9 protein, and 8 µg donor plasmid to the cell solution. H23 cells were electroporated at 200 V with a 45 ms pulse using a square wave protocol.

For B16 mouse melanoma cells the GFP-LaminB1 donor plasmid was designed using the mouse LaminB1 genomic sequence retrieved from NCBI Genome Data Viewer. The genomic sequence was then submitted to AZENTA to construct and subclone into a plasmid vector. The plasmid was transformed in DH5-alpha competent cells to generate plasmid for transfection. B16 cells were transfected using Lipofectamine CRISPRMAX following the manufacturer**’**s guidelines. GFP-positive B16 colonies were sorted using FACS and expanded. Cell sorting was repeated three times over the course of weeks to acquire a pure population of GFP-positive cells.

### Genomic validations of ChReporters

#### Single nucleotide polymorphism array (SNPa)

Genomic DNA was isolated from ∼1 million H23 cells dissociated from spheroids cultured in 1% methylcellulose using the Blood & Cell Culture DNA Mini Kit (Qiagen) following the manufacturer**’**s instructions. DNA samples were sent to the Center for Applied Genomics Core at the Children**’**s Hospital of Philadelphia for analysis using the HumanOmniExpress-24 BeadChip Kit (Illumina), which includes over 700,000 probes spaced approximately 4kb apart across the genome. The core facility provided GenomeStudio files (Illumina), which were used for the chromosome copy number analysis with the cnvPartition plug-in (Illumina).

#### Polymerase Chain Reaction

Genomic DNA was isolated from B16 GFP-LaminB1 cells using the PureLink Genomic DNA Mini Kit (ThermoFisher Scientific) according to the manufacturer**’**s protocol. DNA concentration and purity were assessed using NanoDrop ND-1000 Spectrophotometer. PCR amplification was performed using Platinum II Hot Start Green PCR Master Mix (ThermoFisher Scientific) following the manufacturer**’**s recommended protocol.

The primers used for amplification were:

GFP-Forward: 5**’** CTGCTGCCCGACAACCAC

GFP Reverse: 5**’**TCACGAACTCCAGCAGGAC

Mouse LaminB1 Forward (HA1): 5**’** CTCGAGGCCGACGGTTGC

Mouse LaminB1 Reverse (HA2): 5**’** CCAGCTCGGTCTCGTAGAG

The thermal cycling conditions were 94°C for 2 minutes, followed by 35 cycles of denaturation at 94°C for 15 seconds, annealing at 60°C for 15 seconds and extension at 68°C for 23s. PCR products were analyzed by gel electrophoresis using 1% agarose.

### Immunoblotting

B16 cells were lysed in RIPA lysis buffer (Thermo Fisher Scientific) supplemented with a protease inhibitor (Roche). The Bradford Assay was used to assess protein concentrations, and equal amounts of protein were separated by SDS-PAGE and transferred onto nitrocellulose membranes. The membranes were blocked in 5% non-fat dry milk in TBST for 1 hour at room temperature and incubated overnight at 4C in blocking buffer containing the following primary antibodies: Lamin B1 (Abcam; 1:1000) and GAPDH (Cell Signaling; 1:1000). After washing with TBST, the membranes were incubated with HRP-conjugated anti-rabbit secondary antibody (Cytiva;1:10,000) for 1 hour at room temperature. Protein bands were developed using 3,3**’**,5,5**’**-Tetramethylbenzidine (TMB) (Sigma), scanned and analyzed using Image J.

### Flow Cytometry and Fluorescence activated cell sorting (FACS)

Flow Cytometry analysis was performed using the Penn Cytomics and Cell Sorting Shared Resource Laboratory with H23 GFP-LaminB1, or B16 GFP-LaminB1cells. Gating was performed to select single cells and determine if they were chromosome reporter positive or negative.

Prior to any experiment in which the ChReporter-negative population was assessed, FACS was used to sort H23 GFP-LaminB1, B16 GFP-LaminB1 or A549 RFP-LaminB1 cells (only for 2D) to enrich for the population expressing the chromosome reporter. FACS was performed on BD FACS Aria II.

Flow cytometry analysis of spheroids was performed by first pipetting each spheroid from its well into a 15 mL conical tube, pooling the spheroids of a replicate. These were spun down in a centrifuge for 5 minutes at 300g, the supernate was aspirated, washed with PBS and repeated 2 times. After a final spin down and aspiration, spheroids were resuspended in 0.05% trypsin. Spheroids were incubated in trypsin until spheroids fully disassociated, typically 20-40 minutes. After incubation, disassociated spheroid cells were spun down, washed and resuspended in 0.3 mL of RPMI media supplemented with 10% FBS. The remaining process is the same as 2D cultures.

Replicates for 2D and spheroids were stained with 1:2000 Hoechst and Flow cytometry was performed for 10,000 cells. FACS analysis was performed using FCS express 7 Research Edition.

### Spheroid Cultures

To obtain a given % agarose hydrogels, appropriate amounts of agarose powder were dissolved in PBS (Sigma) at 90°C, then cooled down to 45°C before use.

Freshly-sorted, ChReporter-positive H23 and B16 cells were cultured in 1% (w/v) methylcellulose (Fisher Scientific, Catalog no. S25427) in RPMI 1640, F12 or DMEM media respectively, supplemented with 10% FBS and 1% penicillin-streptomycin. Spheroid cultures in methylcellulose were carried out in 35-mm untreated petri dishes or non-adhesive 96 U-well plates to prevent adhesion of cells and were cultured for 4 days. For spheroid cultures in agarose hydrogels, an initial layer of agarose was cast into a 24-well plate. 50-cell clusters of H23 or B16 cells or 100-cell clusters of A549 or U2OS cells that were formed overnight were then mixed with the agarose hydrogel and placed on top of the initial layer of agarose following the gelation of the initial layer at room temperature. Spheroids were cultured in agarose for 7 days.

### Fluorescence staining & analysis

#### Cancer spheroids

Spheroid samples were fixed using 4% paraformaldehyde in PBS for 45 minutes at room temperature, then washed three times with PBS, blocked with 10% goat serum in 0.5% Triton X-100 in PBS (PBST) for 2 hours, then incubated overnight with anti-rabbit cleaved caspase-3 (Cell Signaling; 1:100). The samples were washed four times at 15 minutes each using 0.5% Triton X-100 in PBS (PBST), then incubated with an appropriate Alexa Fluor secondary antibody (Invitrogen; 1:200). To label F-actin, Alexa Fluor-conjugated phalloidin (Invitrogen, 1:400) was used. Samples were incubated with Hoechst 33342 (1:2000; Invitrogen) for 1 hour at room temperature to label for nuclei and washed three times with PBS for 10 min each time. Spheroids were placed in imaging chambers and incubated in glycerol for a minimum of 1 day, then imaged using the Leica Stellaris laser scanning confocal microscope using a x40 magnification water objective. Fluorescence images were analyzed using Image J.

#### 2D monolayer experiments

H23 and A549 cells were fixed using 4% paraformaldehyde in PBS for 15 minutes at room temperature, then washed three times with PBS, and stained with Hoechst 33342. Wide-field microscopy images were captured at 10x and 40x. Image J and Matlab were used to analyze chromosome loss rate and micronuclei induction following each treatment for both cell lines at a suitable recovery timepoint (not shown).

### Micropipette aspiration

A glass capillary (World Precision Instruments TW100-3) was pulled using a pipette puller (Sutter Instruments P-97). The pulled pipette was then scored using a ceramic tile and bent using a microforge device. To prevent adhesion of spheroids, the glass pipettes were incubated in 1% pluronic F-127 solution in PBS. Spheroids that were cultured in methylcellulose were transferred to a glass-bottom dish. Spheroids were aspirated using a syringe and the pressure difference (ΔP) was measured using a pressure transducer (Validyne). Tissue aspiration (ΔP>0) and retraction (ΔP=0) were recorded using time-lapse microscopy.

### Surface layer and Core cell numbers

Multiple differences between cells in a surface layer on the spheroid relative to cells within the spheroid core motivate some simple estimates. For a first model of small spheroids with relatively few cells, we assume a spheroid of diameter *D* has many smaller spherical cells of diameter *d* (not shown): we show that if *D* > 4*d*, then *N*_core_ > *N*_surf_. For the three spheroids shown in **Figure 1E**, only the smallest spheroid in 2% agarose appears to have fewer core than surface cells, given that *D* ∼ 4*d*. In particular, the cross-section has ∼15 perimeter cells, and ∼13 central cells. These 2D numbers must be corrected for 3D surface and core numbers as:

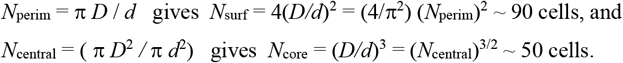

Whereas the above numbers confirm fewer core than surface cells for the small spheroid in 2% agarose, for the one large spheroid in methylcellulose shown in **Figure 1D**:

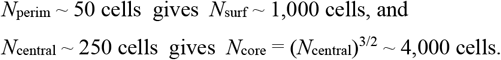

### In vivo tumor studies using B16 GFP-LaminB1 cells

Cells were sorted to obtain a pure GFP+ population. A total of 800,000 cells were injected subcutaneously into C57 mice (n=3) and allowed to form tumors over 19 days. Tumors were isolated and disassociated using dispase, collagenase and DNAase. The tumor cells were stained with Hoechst and an in house anti-ERV stain which labels a retroviral marker on B16 cells to distinguish them from normal cells. Flow cytometry was used to analyze GFP+ and GFP-populations.

A portion of the disassociated cells were cultured in T75 flasks for a week, then fixed with 4% PFA, stained with Hoechst (1:2000), and imaged using an Olympus widefield fluorescent microscope. Remaining cells from the tumor were plated in a 6-well glass bottom plate, fixed after reaching confluency, and similarly stained with Hoechst and anti-ERV before imaging. GFP-negative B16 cells from each tumor were sorted and subjected to PCR analysis.

### Theory extensions of the Luria-Delbruck Model

Data reported in **Figure 2H,I** were modeled through an adaptation of Luria-Delbrück**’**s (LD) theory on random and heritable mutations in bacteria. Starting with their work^20^, we derive a model to describe the mean of *n* cancer cells sampled from a larger population and analyzed using flow cytometry. In LD theory of bacteria mutations, when the mutation rate and sampling is sufficiently low, the mean will usually omit jackpot scenarios, or the time in which a mutation is very unlikely. Hence, it is important to distinguish between two cases. Case 1: Chromosome loss rate, a, is sufficiently high such that one mutation is likely to occur in the first division with a given number of starting cells. Case 2: a is small so it is not likely for a mutation to occur in the first division with a given number of starting cells. We can define a threshold for the two cases:

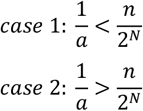

The number of population divisions is represented here by N. These cases point to a threshold where one mutation is likely to occur in the first doubling.

<u>Case 1</u>: We start with Luria-Delbrück**’**s expression for the average number of mutants, ρ, after the exponential growth, *N*_*t*_ = *e*^*t*^:

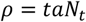

In LD work, time or t is normalized to *t* = In(2)*N*. For the purpose of our experiments, we are interested in setting LD equation as a function of the number of divisions rather than time. This can be rewritten using the following, *N*_*t*_ = *e*^*t*^ = 2^*N*^:

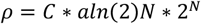

This is the expected average of cells to have lost a chromosome after N divisions for *C* starting number of cells, 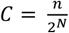.To find the fraction of cells expected to have lost a chromosome after a given number of divisions we simply divide by the total number of cells, *n*, or *C* * 2^*N*^.

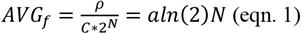

Equation 1 gives us the mean fraction of cells which have lost the chromosome of interest if the mutation rate satisfies case 1 conditions.

<u>Case 2</u>: In this case we wish to omit conditions that are unlikely to occur, just as in LD work we find N_0_ such that 1 mutation or chromosome loss has occurred. If the change in the number of mutated cells, dm, for a small change in division, dN:

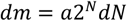

The number of divisions, N_0_, for one mutation is then:

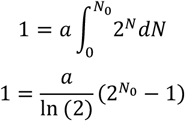

It is important to note that the number of divisions is not much greater than 1. This prevents approximating as 2^No^. We must then solve for starting with *n*/2^*N*^ cells.

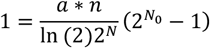

We can solve for N_0_ as a function of N:

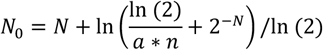

The above can be placed into LD**’**s equation and solved for *C* colonies:

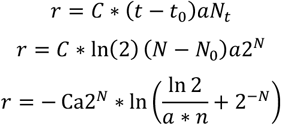

And then a similar process to case one can be applied to find the *AVG*_*f*_:

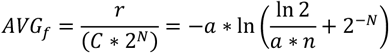

For 10,000 cells:

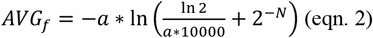

As spheroids typically have a higher fraction of GFP negative, we assume that generally spheroids can be modeled with eqn. 1 and 2D cultures are modeled with eqn. 2. These models are applied to multiple cell types, H23 (**Figure 2H**) and B16 (**Figure 2I**), using MATLAB R2023b and appear to fit the data well. From these we can estimate an upper bound on the chromosome loss rate, which we report in this work as *a*_*u*_.

Although not explored in these experiments, the variance can also be derived starting from Luria-Delbrück**’**s work following a similar method. It should be noted that these experiments are a sampling of a larger population of cells so the Central Limit Theorem would need to be taken into consideration. Time is also not large, so the approximation made by LD does not apply. In addition, further complexity is introduced into deriving the variance when measuring spheroids as cells come in bundles of spheroids, rather than being random in the large 2D cell cultures.

### TCGA and CCLE data mining

Our methods have been recently described ^44,45^. Briefly, for TCGA analysis, expression (RNA-seq, rsem reads), copy number variation (masked cnv), and phenotype data was downloaded from UCSC Xena website (https://xenabrowser.net/datapages/). Measurements of CNV versus LOH are respectively based on amounts of sequence versus specific sequence of alleles, and different methods have different sensitivities for the two types of measurements. LOH can be complex such as a loss followed by duplication of the allele, which yields no CNV but copy-neutral LOH.

For CCLE, CNV (CCLE_segment_cn) and phenotype data was downloaded from the DepMap portal (https://depmap.org/portal/download/all/). CCLE copy number was reported as the ratio between the copy number and the basal reference of the sample, a region with two copies of Chrs will have a ratio of one. The aneuploidy level was obtained by summing |reported ratio – 1| × segment length of the reported ratio within each sample. All aneuploidy levels were normalized to the maximum.

### Statistics

We adhered to standard reproducibility requirements in cell biology such as those outlined by the Journal of Cell Biology. Analysis and model fitting was generally conducted using Prism (Graphpad) and SigmaPlot (SPSS). Model significance was determined using an F-test. Unless indicated, plots display mean ± SEM, with indications for the number of technical replicates (e.g., cells), as well as the number of biological replicates beyond duplicates.

Statistical analyses were conducted using two-tailed Student**’**s t tests when data passed normality tests (e.g. Shapiro–Wilk test in SigmaPlot) and Mann–Whitney U-test was performed if the data did not pass a normality test. When data from multiple experiments were normalized to the control, one-sample t tests were conducted to test for statistical significance. K–S tests were used to compare distributions where relevant. Significance is indicated with a star where appropriate in a figure or legend if P < 0.05.

## Acknowledgements

We thank staff for training and assistance at the Cell & Developmental Biology Microscopy Core (Penn SOM), particularly Drs. Andrea Stout and Jasmine Zhao. We sincerely thank Peter K Zhu and Brandon Hayes for preliminary experiments on H23 cells. This work was supported by funding from the following sources: NIH U01 CA254886 (DED), P01 CA265794 (DED), National Science Foundation CMMI1548571,154857. The authors also acknowledge the additional University of Pennsylvania core facilities: the Penn Cytomics and Cell Sorting Resource Laboratory, and the Penn Genomic and Sequencing Core.

## Author contributions

A Anlas, M Sprenger, and D.E. Discher designed experiments. A Anlas, M Sprenger, N Ontko, and S Phan performed the experiments. A Anlas, M Sprenger, S Phan and D.E. Discher analyzed the data. N Ontko provided mouse tissue for the experiments. A Anlas, M Sprenger, and D.E. Discher wrote the manuscript. Disclosures: The authors declare no competing interests.

## Institutional Review Board Statement

The animal study protocol for mouse B16 tumors was approved by the Institutional Review Board (or Ethics Committee) of the University of Pennsylvania (IACUC-approved protocols #803177 and #804455 with dates of approval of 09/24/2021 and 06/20/2021).

## Informed Consent Statement

Not applicable.

## Data availability

The main data supporting the results in the study are available within the paper and its Supplementary Information. Source data are provided with this paper and are openly available under the author**’**s names at https://figshare.com/. No large sequencing datasets were generated as part of this study, yet publicly available data from The Cancer Genome Atlas were accessed via the identifiers provided in Methods and in the relevant figure captions.

## Code availability

Not applicable.

## Notes

### Competing Interest Statement

The authors have declared no competing interest.

